# Integrative taxonomy reveals hidden species within the western Atlantic *Callichirus major* s. l. (Decapoda, Axiidea, Callichiridae)

**DOI:** 10.1101/2020.09.21.307249

**Authors:** Patricio Hernáez, Marcel S. Miranda, Juliana P. P. Rio, Marcelo A.A. Pinheiro

## Abstract

The ghost shrimp *Callichirus major* (Say, 1818) is widely distributed in the Atlantic Ocean from ∼23°N to ∼26°S, and has also been reported from the tropical eastern Pacific. Evidence has been accumulating over many years that *C. major* is actually a species complex. Yet, the name *C. major* is widely and frequently used in many kinds of research. The current lack of clarity in the use of the name *C. major* has resulted in nomenclatural instability, but also in unreliability and miscommunication of the available ecological and distributional information. Existing morphological and molecular evidence is reviewed and new evidence presented for the specimens from the southern localities previously assigned to *C. major* s. l. actually being a new species. That new species is herein described based on morphological and molecular evidence. Additionally, a neotype is selected for *C. major* in order to settle the defining characters of *C. major* s. str. and, therefore, ensuring the correct use of this name.

## Introduction

The ghost shrimp *Callianassa major* Say, 1818, was based upon a single specimen from the bay shore of the river St. John, east Florida, and subsequently transferred to *Callichirus* by Stimpson (1866) as the type species of his new genus. This species has been recorded over the years from many localities along the western Atlantic coasts where it has a broad distribution ranging from North Carolina (∼23°N) to Santa Catarina (∼26°S), Brazil, including the Gulf of Mexico, and the Caribbean coast of Colombia, but has also been reported from the eastern tropical Pacific (Hay and Shore 1918; Felder 1973; Rodrigues 1983; Williams 1984; Melo 1999; Felder 2001; Sakai 2011).

However, evidence has been accumulating over many years that *C. major* is actually a species complex (Rodrigues and Shimizu 1997; Strasser and Felder 1998, 1999a, b, c; Staton and Felder 1995; Felder and Robles 2009; Peiró 2012). Yet, the name continues to be widely and frequently used in ecological, distributional, morphological, checklists, and taxonomic researches carried out along its implied geographic distribution (Rodrigues 1966, 1971, 1983; Souza and Borzone 1996; Blanco-Rambla 1997; Souza et al. 1998; Coelho et al. 2007; Botter-Carvalho et al. 2007; Botter-Carvalho et al. 2012; Rio et al. 2019; Souza et al. 2020). The current lack of clarity in the use of the name *C. major* has hence resulted in nomenclatural instability, but also in unreliability and miscommunication of the available ecological and distributional information.

The study of a wealth collection of *Callichirus* assembled by the first author along the Brazilian coast from north to south along with the study of males and females from many different localities from the northwestern and southwestern Atlantic in Museum collections, prompted us to review the existing morphological and molecular evidence for the specimens from the southern localities previously assigned to *C. major* actually being a new species. That new species is herein described. Additionally, a neotype is selected for *C. major* (Say, 1818) in order to settle the defining characters of *C. major* s. str. and, therefore, ensuring the correct use of this name.

## Material and methods

Most specimens obtained along the Brazilian coast were collected during the low tide using a yabby pump (diameter = 77 mm, length = 100 cm) in the intertidal zone (< 1 m depth) (Fig. 1a–c). Otherwise, the studied material belongs to collections from the following institutions: MZUSP (Museu de Zoologia, Universidade de São Paulo), USNM (National Museum of Natural History, Smithsonian Institution, Washington D.C.), MZUC (Museo de Zoología, Universidad de Concepción), and MZUCR (Museo de Zoología, Universidad de Costa Rica). The holotype and the paratypes of the new species described herein are deposited at the MZUSP.

**Fig. 1.**
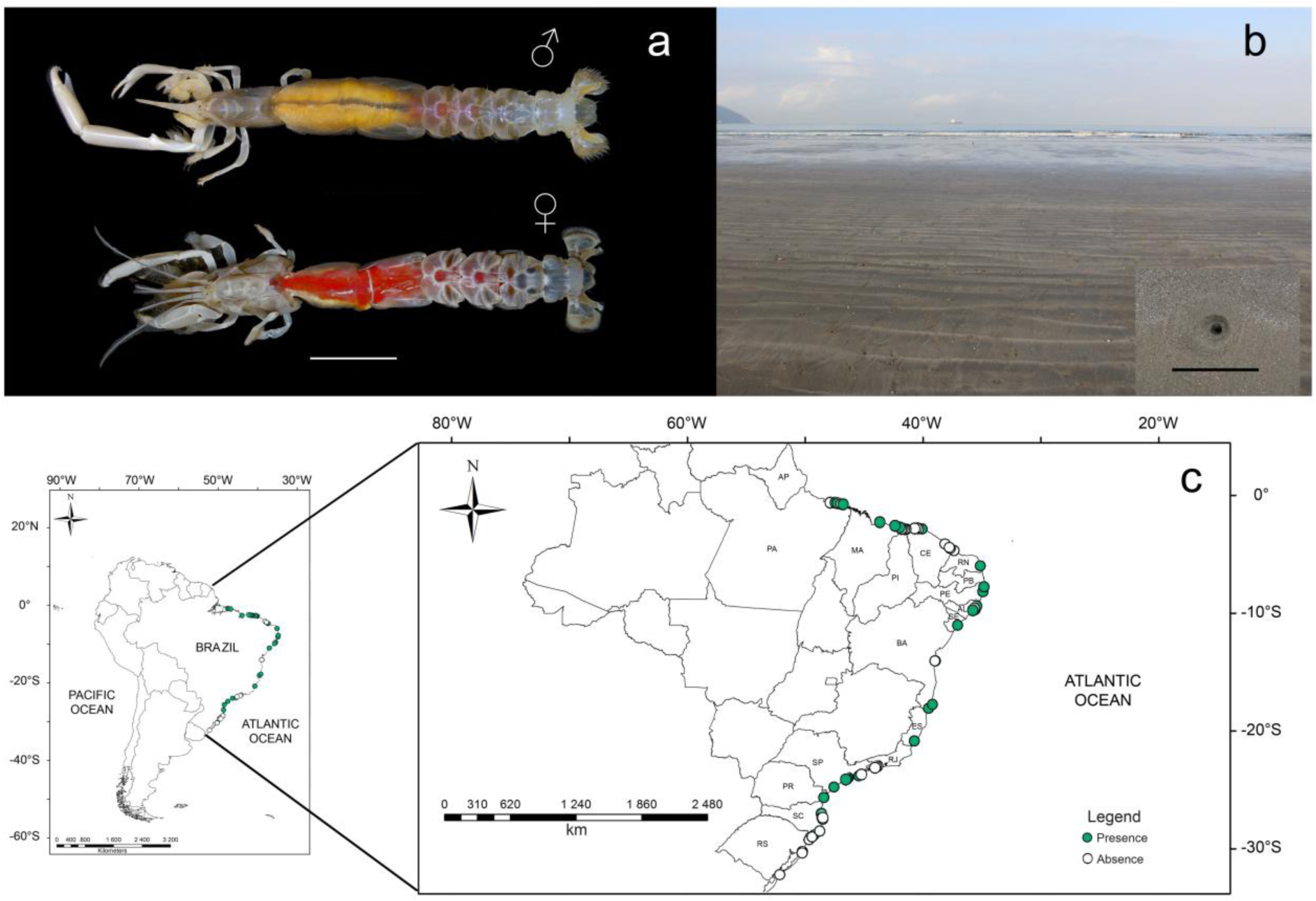
a, male and female specimen of the burrowing shrimp *Callichirus meridionalis* sp. nov. from the Brazilian coast (MZUSP 39027); b, intertidal flat at Gonzaga beach showing a topotypical habitat of the new species, inset indicating a burrow opening of *C. meridionalis* sp. nov.; c, collection sites for *C. meridionalis* sp. nov. along the Brazilian coast. Scale bar: 5mm.

Measurements (mm) were made under a stereomicroscope (Zeiss^®^ Stemi^®^ SV-6) equipped with digital analysis image system (Zeiss^®^ AxioCam^®^ MRc5). Each image was digitized using an electronic tablet for graphic design (Wacom^®^). Size is expressed as postorbital carapace length (cl) measured along the middorsal line of the carapace from the anterior margin of rostrum to the posterior margin of the carapace.

Abbreviations include: Al (antennule or first antenna), A2 (antenna or second antenna), tl (total length), coll. (collector or collected by), Mxp3 (maxilliped 3), Pl (pereopod 1), Plp 1-2 (pleopods 1 and 2), EP (eastern Pacific), WP (western Pacific), EA (eastern Atlantic), WA (western Atlantic).

### Molecular data analysis

The partial sequences of the mitochondrial loci 16S rDNA were retrieved from the GenBank (Table 1). All *Callichirus* species with suitable 16S rDNA sequences available were included. Other Callichiridae and Axiidea species were included to test the monophyletic status of *Callichirus* (Table 1). Sampling locality information, GenBank accession numbers, museum voucher numbers and references for taxa included in the molecular analyses are provided in Table 1.

**Table 1.**
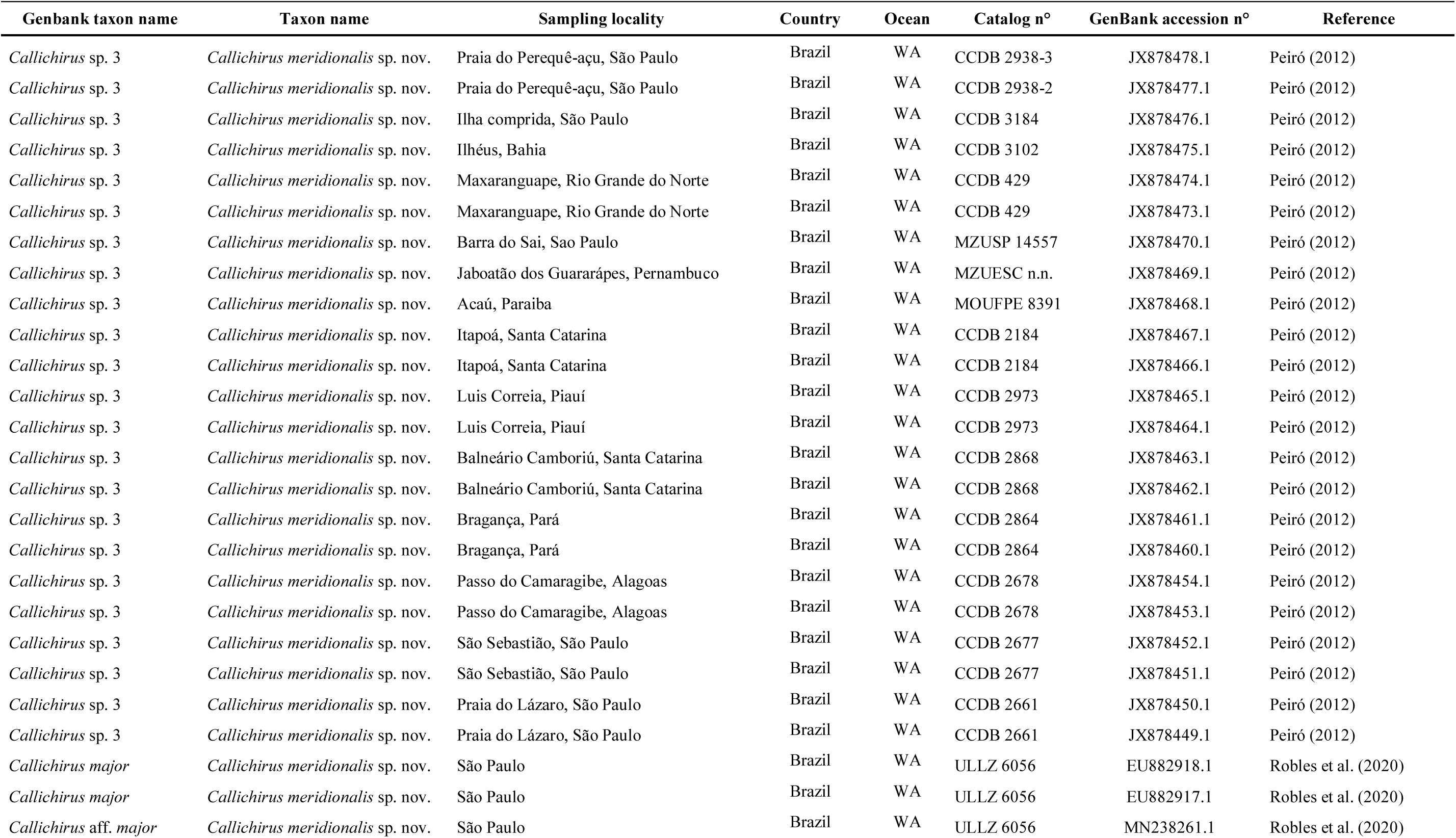
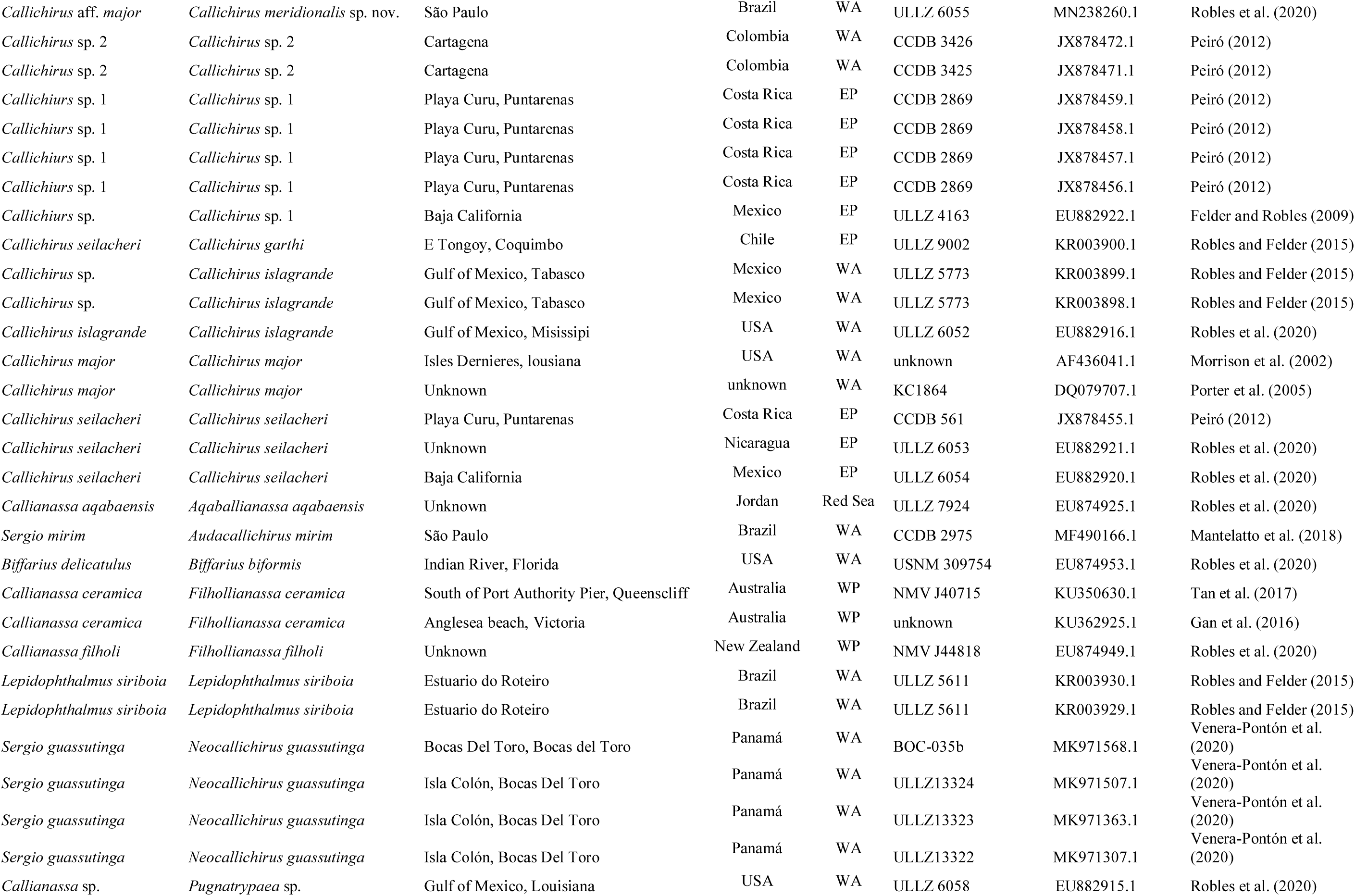
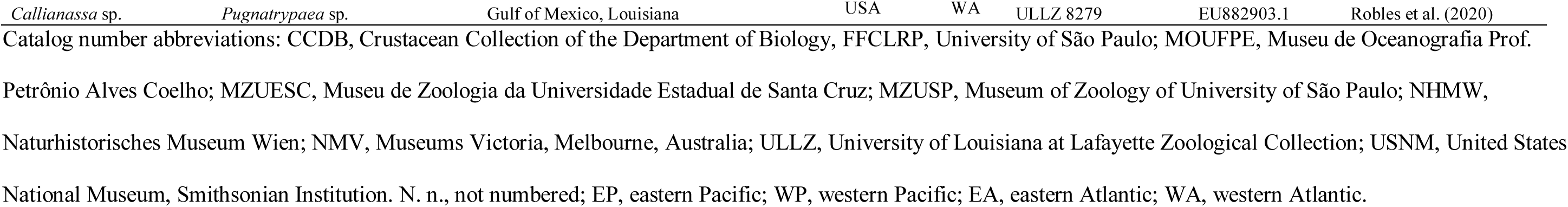
Taxa included in the molecular phylogenetic analyses to place *Callichirus meridionalis* **sp. nov.** within the context of the genus *Callichirus* Stimpson, 1866.

The phylogenetic analysis was inferred by using the Maximum Likelihood method and General Time Reversible model (Nei and Kumar 2000). The tree with the highest log likelihood (−2817.72) is presented. Initial tree(s) for the heuristic search were obtained automatically by applying Neighbor-Join and BioNJ algorithms to the matrix of pairwise distances estimated using the Maximum Composite Likelihood (MCL) approach, and then selecting the topology with superior log likelihood value. A discrete Gamma distribution was used to model evolutionary rate differences among sites (5 categories (+G, parameter = 0.4201)). The rate variation model allowed for some sites to be evolutionarily invariable ([+I], 33.03% sites). The tree is drawn to scale, with branch lengths measured in the number of substitutions per site. This analysis used 54 nucleotide sequences with a total of 455 positions in the final dataset. Phylogenetic analyses were conducted in MEGA X (Kumar et al. 2018). Branch support values were calculated using bootstrap analyses with 1,000 replicates (Felsenstein 1985). Bootstrap support values are shown on nodes of the phylogenetic tree, when support is greater than 50. Nucleotide divergence estimated from pairwise distance was calculated in MEGA X with the same best-fit model (Table 2).

**Table 2.**
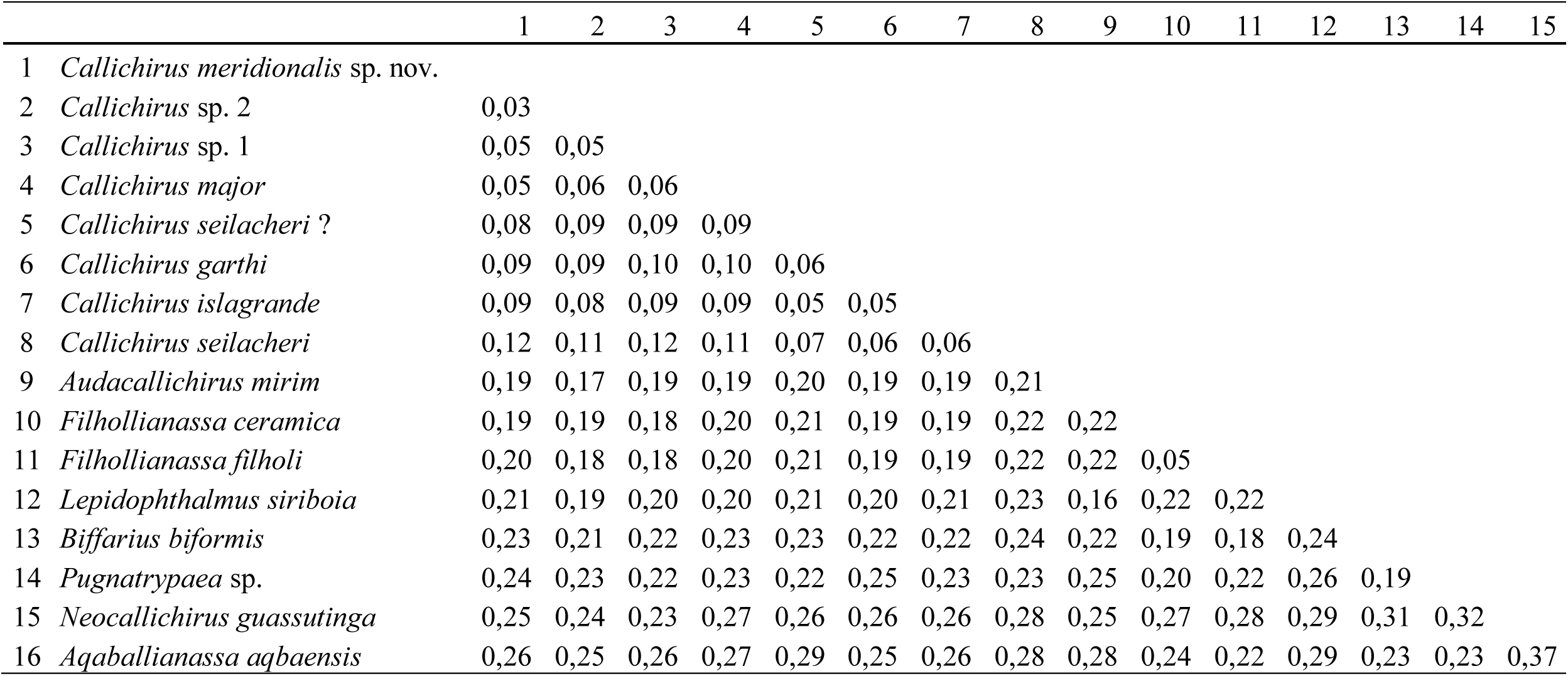
Estimation of evolutionary divergence over sequence pairs from the portion of the mitochondrial 16S rDNA with ∼510bp.

## Results

### Taxonomy

**Infraorder AXIIDEA de Saint Laurent, 1979**

**Family CALLICHIRIDAE Manning & Felder, 1991**

**Genus *Callichirus* Stimpson, 1866**

#### Type species

*Callichirus major* (Say, 1818), by original designation [Type locality: Bay shore of the river St. John, east Florida, USA].

**Included species** (genera of the original combination indicated within brackets) **–** *Callichirus adamas* (Kensley, 1974) [*Callianassa*] (EA); *C. garthi* (Retamal, 1975) [*Callianassa*] (EP); *C. islagrande* (Schmitt, 1935) (WA); *C. meridionalis* sp. nov. (WA); *C. major* (Say, 1818) [*Callianassa*] (WA); *C. santarosaensis* Sakai & Türkay, 2012 (WA); and *C. seilacheri* (Bott, 1955) [*Callianassa*] (EP).

***Callichirus meridionalis* sp. nov**. (Figures 1a, 2, 3, 4)

**Fig. 2.**
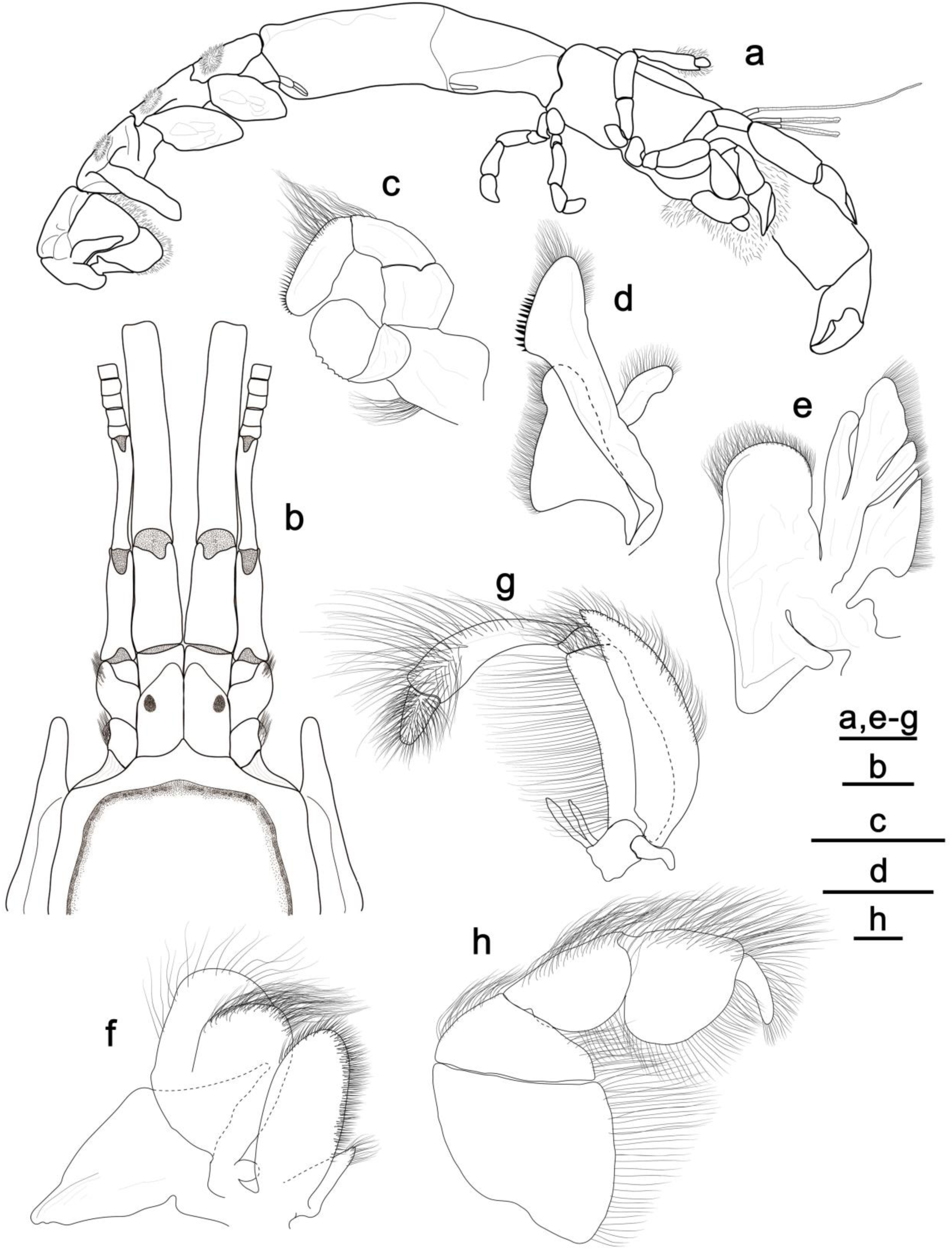
*Callichirus meridionalis* sp. nov. a, b, holotype, male, cl 18.8 mm, MZUSP 41251; c– h, paratype, male, cl 18.6 mm, MZUSP 41253. a, body, lateral view; b, carapace front, eyestalks, and antennular and antennal peduncles, dorsal view; c, right mandible, external surface; d, right maxillulla, external surface; e, right maxilla, external surface; f, right first maxilliped, external surface; g, right second maxilliped, external surface; h, right third maxilliped, external surface. Scale bars: a = 1 cm; b–h = 2 mm.

**Fig. 3.**
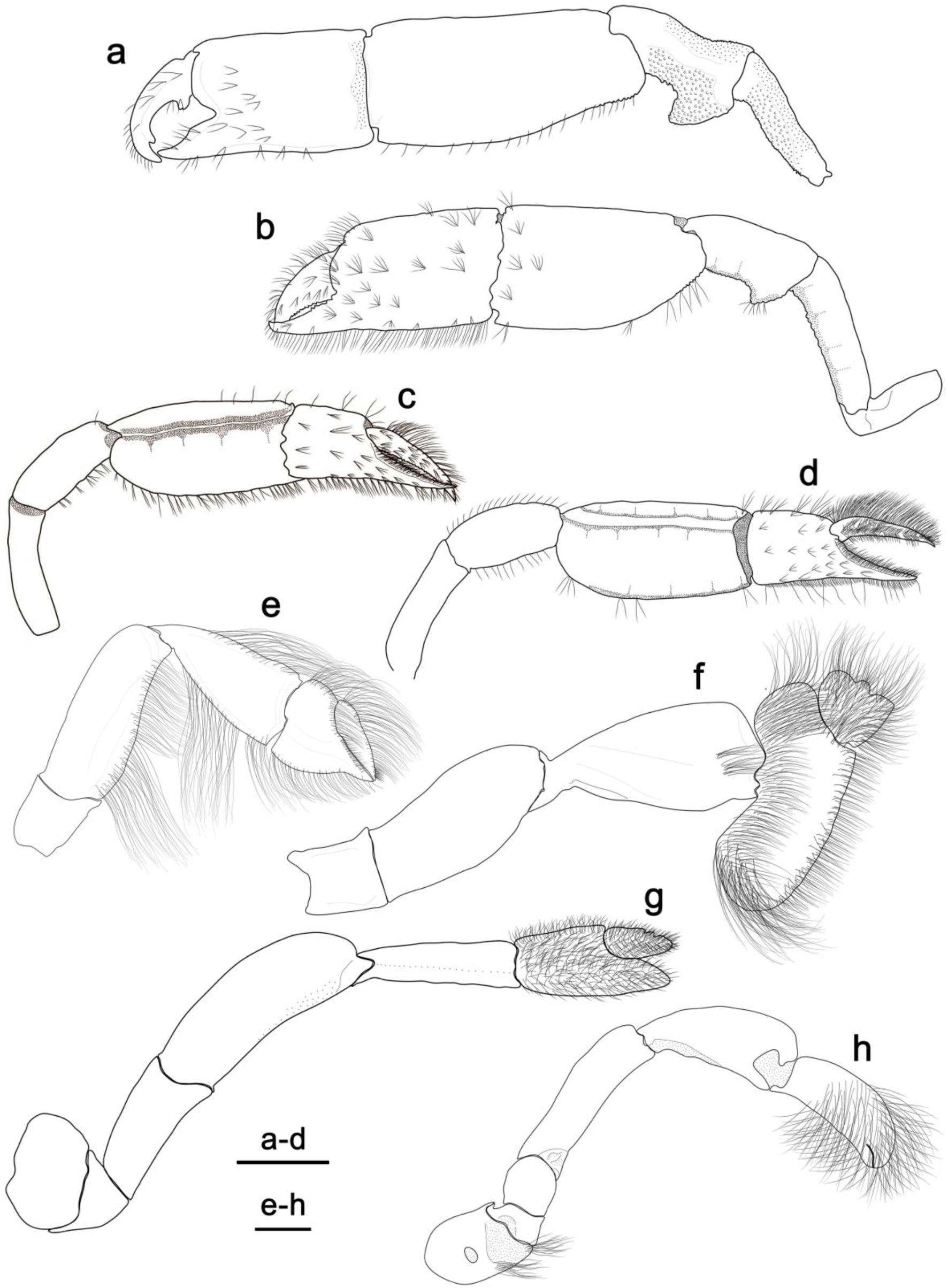
*Callichirus meridionalis* sp. nov., a, c, e–h, holotype, male, cl 18.8 mm, MZUSP 41251; b, d, paratype, female, cl 20.4 mm, MZUSP 41252. a, c, male major and minor chelipeds, respectively, lateral view; b, d, female major and minor chelipeds, respectively, lateral view; e–h, second, third, fourth, and fifth pereopods, respectively, lateral view. Scale bars: a–d = 1 cm; e–h = 2 mm.

**Fig. 4.**
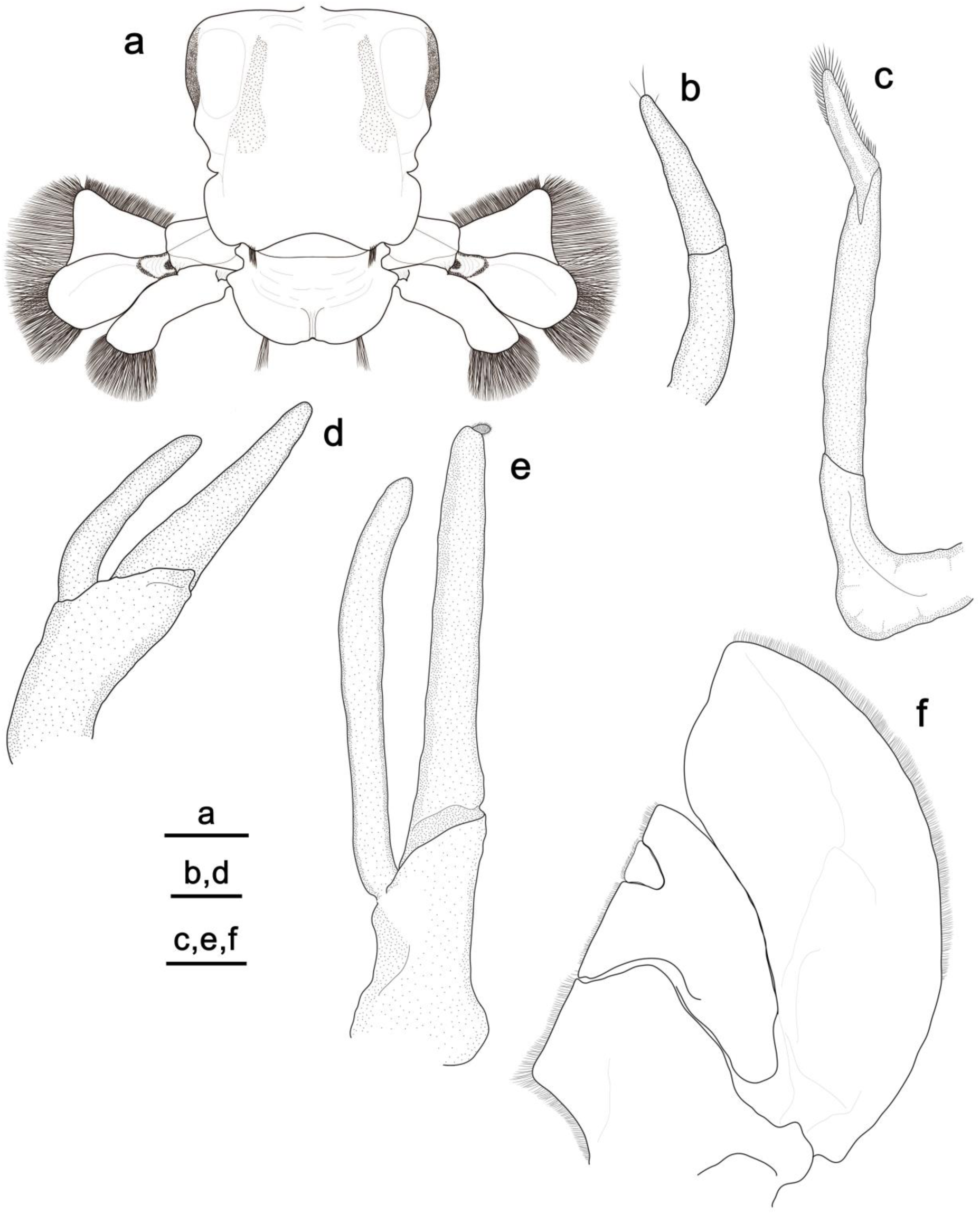
*Callichirus meridionalis* sp. nov., a, b, d, f, holotype, male, cl 18.8 mm, MZUSP 41251; c, e, paratype, female, cl 20.4 mm, MZUSP 41252. a, sixth abdominal somite, uropods, and telson, dorsal view; b, c, male and female first pleopods, respectively, external surface; d, e, male and female second pleopods, respectively, external surface; f, third to fifth pleopods, external surface. Scale bars: a, c, e, f = 2 mm; b, d = 1 mm.

*Callianassa* (*Callichirus*) *major –* Rodrigues 1971: 192, figs 1-20 [not *Callichirus major* (Say, 1818)].

*Callichirus major* – Rodrigues 1976: 85, figs. 1-35; 1985: 195, figs. 1-30; Rodrigues and Höld 1990: 48; Borzone and Souza 1996: 67; Souza and Borzone 1996: 553; Rodrigues and Shimizu 1997: 155, fig. 1; Souza et al. 1998: 151; Coelho and Rodrigues 2001: 1447, figs. 13-21; Botter-Carvalho et al. 2002: 97; Souza and Borzone 2003: 625; Botter-Carvalho et al. 2007: 508; Peiró and Mantelatto 2011: 5; Peiró et al. 2011: 261; Botter-Carvalho et al. 2012: 89; Dworschak et al. 2012: 151, fig. 69.29a,b, 69.31t; Alves-Junior et al. 2014a: 109; Alves-Junior et al. 2014b: 13; Alves-Junior et al. 2018: 166; Peiró et al. 2014: 294; Peiró and Mantelatto 2016: 103, pl. 1; Hernáez et al. 2018: 97; Rosa et al. 2018: 1; Hernáez et al. 2019: 1, fig. 1; Rio et al. 2019: 1, figs. 1A, 2; Hernáez et al. 2020: 1, fig. 4c; Moschetto et al. 2020: 1; Laurino et al. 2020: 1 [not *Callichirus major* (Say, 1818)].

*Callichirus* sp. – Strasser and Felder 1999a: 865.

*Callichirus macrotelsonis* – Peiró, 2012: 58, figs. 3-7 [*nomen nudum*].

*Callichirus brasiliensis* Rio, 2018: 20, figs. 1A, 2A, 3A, 3B, 4A [*nomen nudum*].

#### Type material

Brazil: São Paulo: holotype, male, cl 18.8 mm (MZUSP 41251), Praia do Gonzaga, 23°58’13”S, 46°20’04”W, lower intertidal, Santos, São Paulo, P. Hernáez coll., 1 September 2016. Paratypes: 1 female, cl 20.4 mm (MZUSP 41252), 4 males (one dissected), cl: 15.4–19.4 mm, and 4 females, cl 16.4–22.2 mm (2 ovigerous females) (MZUSP 41253), same data as holotype.

#### Non type material

Brazil: Pará: 8 males, cl: 10.1–18.3 mm and 15 females, cl: 10.4–19.2 mm, Praia do Crispim, 00°34’58”S, 47°39’04”W, lower intertidal, Marapanim, coll. P. Hernáez, 9 July 2017 (MZUSP 38995); 8 males, cl: 13.7–19.6 mm and 11 females, cl: 10.5–21.2 mm, Praia de Ajuruteua, 00°49’40”S, 46°36’20”W, lower intertidal, Bragança, P. Hernáez coll., 7 July 2017 (MZUSP 38993). Maranhão: 10 males, cl: 10.8–17.6 mm and 15 females, cl: 12.5–19.8 mm, Praia Olho d’agua, 02°28’44”S, 44°13’51”W, lower intertidal, São Luis, P. Hernáez coll., 6 July 2017 (MZUSP 38994); male, cl 12.4 mm and 2 females, cl: 15.6–18.7 mm, Praia de Tutoia, 02°45’40”S, 42°15’45”W, lower intertidal, Tutóia, P. Hernáez coll., 5 July 2017 (MZUSP 39001). Piauí: 9 males, cl: 9.6–15.6 mm and 14 females, cl: 10.2–17.7 mm, Praia da Atalaia, 02°53’22”S, 41°37’41”W, lower intertidal, Luis Correia, P. Hernáez coll., 3 July 2017 (MZUSP 38998). Ceará: 2 females, cl: 5.8–10.4 mm, close to the marine lighthouse, 02°52’35”S, 40°55’24”W, lower intertidal, Camocim, P. Hernáez coll., 3 July 2017 (MZUSP 39006). Rio Grande do Norte: 22 males, cl: 5.6–12.8 mm and 19 females, cl: 7.6–13.3 mm, Praia do Pirangi, 05°58’28”S, 35°07’28”W, lower intertidal, Pirangi, P. Hernáez coll., 11 June 2016 (MZUSP 39011). Pernambuco: male, cl 10.5 mm and 4 females, cl: 13.9–16.1 mm, Praia de Baixa Verde, 07°45’17”S, 34°49’28”W, lower intertidal, Ilha de Itamaracá, P. Hernáez coll., 17 June 2016 (MZUSP 39012); 6 males, cl: 7.9–15.9 mm and 5 females, cl: 7.8–15.3 mm, Praia Piedade, 08°10’01”S, 34°54’47”W, lower intertidal, Recife, P. Hernáez coll., 10 June 2016 (MZUSP 39013). Alagoas: 15 males, cl: 9.0–16.0 mm and 14 females, cl: 12.9–17.7 mm, Praia do Sobral, 09°40’22”S, 35°33’44”W, lower intertidal, Maceió, P. Hernáez coll., 9 June 2016 (MZUSP 39014). Sergipe: 9 males, cl: 11.5–15.8 mm and 18 females, cl: 10.3–18.0 mm, Praia Aruana, 11°00’50”S, 37°03’50”W, lower intertidal, Aracajú, P. Hernáez coll., 8 June 2016 (MZUSP 39016). Bahia: 16 males, cl: 12.6–15.7 mm and 16 females, cl: 11.8–14.7 mm, Praia Malvinas, 18°04’48”S, 39°32’35”W, lower intertidal, Mucuri, P. Hernáez coll., 20 June 2016 (MZUSP 39022). Espirito Santo: 15 males, cl: 7.8–12.5 mm and 11 females, cl: 6.8–12.2 mm), Praia de Piúma, 20°50’37”S, 40°44’02”W, lower intertidal, Piúma, P. Hernáez coll., 3 June 2016 (MZUSP 39023). São Paulo: 14 males, cl: 5.7–11.9 mm and 08 females, cl: 7.4–11.5 mm, Praia de Barequeçaba, 23°49’39”S, 45°26’04”W, lower intertidal, Barequeçaba, São Paulo, P. Hernáez coll., 2 June 2016 (MZUSP 39028); 28 males, cl: 10.1–19.1 mm and 64 females, cl: 8.0–23.2 mm, Praia do Gonzaga, 23°58’13”S, 46°20’04”W, lower intertidal, Santos, P. Hernáez coll., 1 September 2016 (MZUSP 39027); 3 males, cl: 8.6–15.5 mm and 3 females, cl: 7.9–8.3 mm, Cibratel, 24°12’04”S, 46°48’45”W, lower intertidal, Itanhaém, P. Hernáez coll., 12 August 2016 (MZUSP 39026); 3 males, cl: 7.3–10.8 mm and 3 females, cl: 7.4–14.4 mm, Rio Peruíbe, 24°19’47”S, 46°59’59”W, estuary, Peruíbe, P. Hernáez coll., 1 September 2016 (MZUSP 39025); 8 males, cl: 6.4–18.7 mm and 4 females, cl: 12.6–15.9 mm, Praia Ilha Comprida, 24°45’22”S, 47°33’34”W, lower intertidal, Ilha Comprida, P. Hernáez & J. Rio coll., 1 July 2016 (MZUSP 39029). Paraná: 15 males, cl: 7.9–16.7 mm and 22 females, cl: 10.0–16.4 mm, Praia de Leste, 25°37’50”S, 48°25’16”W, lower intertidal, Pontal de Paraná, P. Hernáez & J. Rio coll., 2 July 2016 (MZUSP 39037). Santa Catarina: 10 males, cl: 11.0–16.7 mm and 27 females, cl: 8.9–22.3 mm, Balneário Camboriú, 26°59’20”S, 48°37’45”W, lower intertidal, Camboriú, P. Hernáez & J. Rio coll., 3 July 2016 (MZUSP 39039).

#### Comparative material examined

*Callichirus garthi* (Retamal, 1975): Chile: holotype, male, cl 35 mm, Lenga, 36°45’S, 73°10’W, Concepción, M.A. Retamal coll., May 1975 (MZUC-UCCC 7311); paratype, male, cl 30 mm, same site as holotype, March 1974, coll. M.A. Retamal (MZUC-UCCC 7313); 66 males, cl: 7.6–21.3 mm and 68 females, cl: 8.6–20.0 mm, Lenga, 36°46’00”S, 73°10’18”W, Concepción, P. Hernáez coll., 9 March 2011 (MUAPCD 0432/2011). *Callichirus islagrande* (Schmitt, 1935): USA: Gulf of Mexico: holotype, male, cl 19 mm, Grand Isle, Louisiana, W.W. Anderson coll., summer of 1930 (USNM 69362). *Callichirus major* (Say, 1818): USA: North Caroline: male, cl 20.6 mm, Onslow Bay, 34°36’23”N, 77°12’35”W, New River, R.B. Manning & D.B. Bixler coll., 30 July 1989 (USNM 266227); female, cl 11.9 mm, Onslow Bay, 34°36’23”N, 77°12’35”W, New River, R.B. Manning & D.B. Bixler coll., 7 August 1990 (USNM 266232). Georgia: male, cl 21.2 mm, Tybee Island, 31°59’58”N, 80°50’29”W, G.A. Bishop coll., 1 April 1988 (USNM 266247). Louisiana: 2 males, cl: 15.0– 24.4 mm, Grand Island, 30°08’N, 89°25’W, Gulf of Mexico, E.R. Willis coll., 26 June 1939 (USNM 79171). Texas: ovigerous female, cl 16.4 mm, Heald Bank, 29°43’36”N, 93°42’29”W, Sabine, W.G. Hewatt coll., 1967 (USNM 97653). Florida: 2 ovigerous females, cl: 9.7–20.2 mm, Indian River, 27°52’48”N, 80°27’24”W, R.B. Manning, W. Lee, M. Schotte & C. King coll., 20 April 1988 (USNM 266125); female, cl 21.0 mm, Indian River, 27°38’11”N, 80°21’50”W, North Hutchinson Island, Fort Pierce, D.L. Felder & W. Lee coll., 14 August 1987, USNM 266118; male, cl 11.1 mm, Fort Pierce area, 27°28’18”N, 80°17’48”W, R.B. Manning coll., 2 March 1987 (USNM 266127); female, cl 15.4 mm, Indian River, 27°28’18”N, 87°17’48”W, North Hutchinson Island, Fort Pierce, R.B. Manning & W. Lee coll., 2 March 1987 (USNM 266126); female, cl 16.3 mm, Indian River, 27°11’00”N, 80°09’30”W, Seminole Shores, R.B. Manning & L.K. Manning coll., 7 July 1984 (USNM 266111); female, cl 16.3 mm, Indian River, 27°10’30”N, 80°10’24”W, Flat Just Inside Saint Lucie Inlet, R.B. Manning coll., 11 February 1983 (USNM 228087); topotypic, male, cl 13.2 mm, Indian River, 27°10’14”N, 80°10’24”W, R.B. Manning coll., 9 February 1983 (USNM 228086); male, cl 18.6 mm, Lake Worth Inlet, 26°46’17”N, 80°02’14”W, Peanut Island, R.B. Manning & D.L. Felder coll., 11 August 1987 (USNM 266114). *Callichirus* aff. *major*: Colombia: Caribbean coast: male, cl 22.2 mm, La Boquilla, 10°28’19”N, 75°29’59”W, Cartagena, R. Lemaitre coll., 10 August 1980 (USNM 266208); male, cl 13.9 mm and ovigerous female, cl 13.7 mm, La Boquilla, 10°28’19”N, 75°29’59”W, Cartagena, R. Lemaitre coll., 10 August 1988 (USNM 266225); female, cl 23.6 mm, Castillo Grande, 10°23’43”N, 75°33’02”W, Cartagena Bay, Cartagena, R. Lemaitre coll., 7 July 1988 (USNM 266211). *Callichirus seilacheri* (Bott, 1955): El Salvador: male, cl 16.3 mm and female, cl 24.1 mm, Los Blancos, topotypic, 13°19’38”N, 88°58’10ʹ”W, P. Hernáez & A. Gamboa-González coll., 22 July 2013 (MZUCR 3335–01); male, cl 8.4 mm and 5 females, cl: 9.7–20.2 mm, Los Blancos, topotypic, 13°19’38”N, 88°58’10”W, P. Hernáez & A. Gamboa-González coll., 22 July 2013 (MZUCR 3336–01). Costa Rica: Pacific coast: 15 males, cl: 8.2–14.5 mm and 16 females, cl: 8.6–14.9 mm, Mata de Limon, 09°55’12”N, 84°42’37”W, Puntarenas, P. Hernáez & A. Gamboa-González coll., 10 June 2012 (MZUCR 3337–01).

#### Diagnosis

Carapace with small triangular rostrum and two anterolateral rounded projections. Ocular peduncle with obtuse tip, distal margin not reaching to second article of antennular peduncle. A1 peduncle stouter, longer than A2 peduncle; second article of antennular peduncle slightly exceeding fourth article of antennal peduncle. Maxilliped 3 merus with distal and proximal margins not parallel, strongly oblique distally, not projecting beyond carpo-meral articulation. Male major cheliped with prominent hook on flexor margin of merus; fixed finger with small triangular tooth on midlength of cutting edge; dactylus strongly arched with tip curved downward and bifid, longer than fixed finger, cutting edge with large bifid tooth proximally, otherwise unarmed. Male pleopod 2 with endopod and exopod well developed. Telson slightly broader than long, tapering distally, emarginate posteriorly.

#### Description

Carapace length of adults up to 22 mm. Carapace smooth, dorsally oval, cervical groove and linea thalassinica distinct, cardiac prominence absent, lateral margin emarginate on central part; rostrum small, short and triangular, rounded anterolateral projections on each side (Figs. 2a, b). Branchial formula as shown in Table 3.

**Table 3.** Branchial formula for *Callichirus meridionalis* **sp**. **nov**.

|  | Maxillipeds |  |  | Pereopods |  |  |  |  |
| --- | --- | --- | --- | --- | --- | --- | --- | --- |
|  | 1 | 2 | 3 | 1 | 2 | 3 | 4 | 5 |
| Exopods | 1 | 1 | - | - | - | - | - | - |
| Epipods | 1 | 1 | - | - | - | - | - | - |
| Podobranchs | - | - | - | - | - | - | - | - |
| Arthrobranchs |  | 1 | 2 | 2 | 2 | 2 | 2 | - |
| Pleurobranchs | - | - | - | - | - | - | - | - |

Ocular peduncle (Fig. 2b) with obtuse tip, slightly separated distally, distal margin not reaching second article of antennular peduncle (A1). Cornea subterminal, darkly pigmented and located distolaterally. A1 peduncle stouter, longer than A2 peduncle; lower margin of antennular peduncle setose. Second article of antennular peduncle slightly longer than fourth article, distal article about twice as long as penultimate article; flagellum with dorsal and ventral ramus of similar length, each one with 21-23 smaller articles, both ramus with sparse, long tufts of setae. A2 peduncle with distal and penultimate article of similar length; basal article with excretory pore protruded; second article with tuft of setae distolaterally; third article shorter than second; fourth article with sparse and long setae, fifth article with tuft of long setae distolaterally, scaphocerite reduced.

Mandible (Fig. 2c) with molar process developed, consisting of one calcareous blunt tooth projected upward; incisor process weakly calcified, distal margin armed with seven acute corneous teeth; palp trisegmented, terminal article longest, external surface covered with long setae from two-third to articulation with penultimate article, with dense, short, stiff setae at distal third. Maxillulle (Fig. 2d) with narrow palp (endopod), terminal article deflected proximally at articulation; proximal endite kidney-shaped, external margin with row of short setae; distal endite elongate, hatchet-shaped, distal margin with short, sharp stiff setae at proximal half. Maxilla (Fig. 2e) foliaceous; proximal and distal endite deeply bilobed, external surface of both heavily setose; endopod narrowing terminally, slightly flexed distally; exopod forming large, broad, scaphognathite.

Maxilliped 1 (Fig. 2f) foliaceous; proximal endite reduced, triangular, setose distally; distal endite developed, subrectangular, external surface heavily setose; endopod rounded, reduced; exopod large, with marked notch on mesial margin, setose marginally; epipod large, with posterior and anterior lobes developed, both narrowing terminally. Maxilliped 2 (Fig. 2g) with long, narrow endopod; dactylus short, less than half of propodus, with terminal brush of short, thick, stiff setae; propodus less than half of merus, strongly arcuate; carpus subtriangular, short; merus more than five times as long as broad, flexor margin with dense fringe of long, simple setae; exopod sword shaped, reaching distal margin of carpus, lateral margin setose; epipod narrow, strongly reduced. Maxilliped 3 (Fig. 2h) without exopod and epipod; ischium-merus operculiform, length about 1.2 times its width, marginally setose; ischium without crest or teeth on mesial surface, 1.4 times length of merus; distal and proximal margin of merus not parallel, strongly oblique distally, not projecting beyond carpo-meral articulation; carpus and propodus expanded ventrally; dactylus digitiform, shorter than propodus.

Chelipeds (pereopods 1) unequal and dissimilar in males and females (Figs. 3a, b). Male major cheliped (Fig. 3a) extremely elongated and strongly calcified; ventral margin of ischium with row of well-spaced blunt teeth along surface, dorsal margin slightly straight; merus with prominent hook on flexor margin, margin of hook strongly serrate, acute distally, remainder of ventral margin with irregular rounded denticles, dorsal margin slightly denticulate and concave; carpus length about 1.5 times than palm, carpus about two times longer than wide, dorsal margin straight, unarmed, ventral margin slightly expanded medially, with rounded denticles at proximal third, sparse setae; palm rectangular, longer than width, dorsal and ventral margins smooth, with sparse setae on ventral margin; fixed finger with triangular tooth on midlength of cutting edge; dactylus strongly arcuate with tip curved downward, bifid, longer than fixed finger, cutting edge with large bifid tooth proximally, otherwise unarmed. Male minor cheliped (Fig. 3c) slender; ischium longer than merus, dorsal and ventral margin smooth; merus narrower than carpus, dorsal margin smooth, slightly curved, ventral margin with spaced tufts of short setae; carpus longest, lateral sulcus, with dorsal margin slightly curved, ventral margin slightly expanded, with spaced tufts of short setae; length of palm about half of carpus length, dorsal margin unarmed, ventral margin with spaced tufts of setae, fixed finger approximately as long as dactylus, cutting edge minutely pectinate; dactylus slightly arcuate, minutely pectinate, with tuft of setae dorsally.

Female major cheliped (Fig. 3b) not as long as in males, differing from male larger cheliped as follows: dorsal and ventral margin of ischium straight; merus with less prominent hook on flexor margin, margin of hook weakly serrate, dorsal margin smooth, curved; carpus as long as palm; cutting edge of fixed finger with two acute teeth at proximal third, as long as dactylus; dactylus slightly curved, cutting edge with small acute teeth. Female minor cheliped (Fig. 3d) as in males, differing only in having a dorsal and ventral margin of merus curved, with cutting edge of fixed finger and dactylus unarmed.

Pereopod 2 (Fig. 3e) chelate, densely setose ventrally; length of ischium about 1.4 times its width; merus slightly longer than carpus, proximo-ventral expansion with long setae, dorsal margin slightly curved; carpus subtriangular, widening distally, dorsal and ventral margin with sparsely long setae; palm less than half of carpus, fixed finger subtriangular, cutting edge smooth; dactylus lanceolate, densely setose dorsally. Pereopod 3 (Fig. 3f) pediform; ischium subsquare, unarmed; merus about two times longer than ischium, ventral margin setose, with weak expansion on central region, dorsal margin non-setose; carpus subtriangular, widening distally, margins setose; propodus expanded ventrally, keel-like, setose, length of dorsal margin shorter than ventral margin; dactylus broad, with two small notches on upper border, surface densely covered with setae. Pereopod 4 (Fig. 3g) not chelate; ischium more than half of merus, unarmed, margins non-setose; merus slightly wider than ischium, arcuate ventrally; carpus conical distally, almost equal in length with propodus, margins non-setose; propodus wider than carpus, surface densely covered with short setae; dactylus reduced, digitiform, with two small notches on the upper border, surface densely covered with short setae. Pereopod 5 (Fig. 3h) chelate, fingers not gaping; ischium rectangular, unarmed; merus longest, about 2.7 times longer than ischium, unarmed; carpus subtriangular, widening distally, unarmed; inner and outer surface of propodus densely setose in distal half; surface of dactylus covered with dense setae.

Pleon (Fig. 2a) smooth, glabrous dorsally except for subcircular patches of setae on lateral margins of somites 3 to 5 representing integumental glands. Somite 2 longest; somite 3-5 similar in length. Somite 6 saddle-like, longer than telson. Telson (Fig. 4a) slightly broader than long, tapering distally, emarginate posteriorly; lateral margin with small lobe proximally, followed by deep notch and another large lobe at midlength to lateral margin; dorsal surface separated into three lobes, one large anterior and two smaller posterolateral separated by deep central notch.

Pleopod 1 (Figs. 4b, c) uniramous in both sexes; two-segmented in males, tri-segmented in females. Pleopod 2 (Figs. 4d, e) biramous in both sexes; in males and females endopod and exopod well developed, exopod slightly shorter than endopod; in males endopod without appendices interna and masculina; in females endopod with well-developed and terminal appendix interna, surface of appendix interna covered with patch of short hooked setae. Pleopods 3 to 5 (Fig. 4f) biramous, leaf-like, endopod broadened, with subtriangular appendix interna embedded into mesial margin of endopod in both sexes. Uropod (Fig. 4a) with protopod dorsally divided into four irregular lobes, posterolateral lobe ending in spinous process projected distally; endopod strap-shaped, much longer than wide, exceeding posterior margin of telson, base with rounded lobe ending in a spine protruded distally; exopod triangular with anterodorsal plate shorter than posterodorsal plate, both with distal margin densely setose.

#### Distribution

Known from Praia do Crispim, Pará to Balneário Camboriú, Santa Catarina, Brazil (Fig. 1c).

#### Habitat

*Callichirus meridionalis* sp. nov. builds simple burrows with one opening (Fig. 1b), normally inhabited by only one individual, in the intertidal zone of fine-grain size sandy beaches. This species often is the only axiidean species present in the intertidal zone of many beaches. The pinnotherid crabs *Austinixa aidae* (Righi, 1967) and *A. patagoniensis* (Rathbun, 1918), are normally found living within of the galleries of *C. meridionalis* sp. nov. (Hernáez 2018).

#### Color

Carapace transparent and hyaline; chelipeds white; first and second pleomeres transparent; deep yellow hepatopancreas and reddish-orange ovary visible in pleonal region of mature females; posterior region of the gonads one has an orange coloration in mature males, denoting an ovarian part of testis and consequently the hermaphroditism of individuals belonging to this sex; telson white and hyaline. Pattern of dorsal abdominal grooves more whitish in adult males and females (Fig. 1a).

#### Etymology

From the Latin, *meridionalis* referring to the southern distribution of the new species.

## Discussion

### Taxonomic background of Callichirus major (Say, 1818)

The identity of *Callichirus major* (Say, 1818) has been shrouded in uncertainty for many years, despite that larval and molecular evidence have been accumulating to suggest that it is in fact a species complex (e.g., Staton and Felder 1995; Strasser and Felder 1998, 1999a; Felder and Robles 2009; Peiró 2012; Rio 2018). Long before molecular data was available to suggest splitting the southern Atlantic population previously assigned to *C. major* into a new species, comparative morphological studies by Rodrigues (1985) (see also Rodrigues 1971) have shown that southern (Santos, Brazil) and northern specimens (North Carolina, USA) differed from each other. The differences concerned the presence of a subterminal tooth on the cutting edge of the dactylus of the male major cheliped (absent in northern specimens) and the absence of an acute tooth at the proximal third of the propodus of the female cheliped (present in northern specimens). Rodrigues (1985) suggested that the Brazilian population of *Callichirus* would ultimately prove to be a different species from *C. major*. Rodrigues felt that more specimens from both northern and southern localities were required before a new species was described. Many years later, Strasser and Felder (1999a) commented on several morphological differences between larvae from northern and southern (Brazilian) localities and referred to the Brazilian material as *Callichirus* sp.

Ortmann (1893: 87) described *Anomalocaris macrotelsonis* from northern Brazil based on a fourth larval stage and commented on the resemblance of *A. macrotelsonis* to the mysis of *Callianassa*. However, Ortmann’s species has a distinct projecting dorsal carina on the first abdominal segment (Ortmann 1893: 87, pl. VI, fig. 5), a characteristic present neither in *Callianassa* and *Callichirus* nor in other callichirid or callianassid larvae known to date (see also Pohle et al. 2011 and Pohle and Santana 2014 for a compilation of available larval information).

**Fig. 5.**
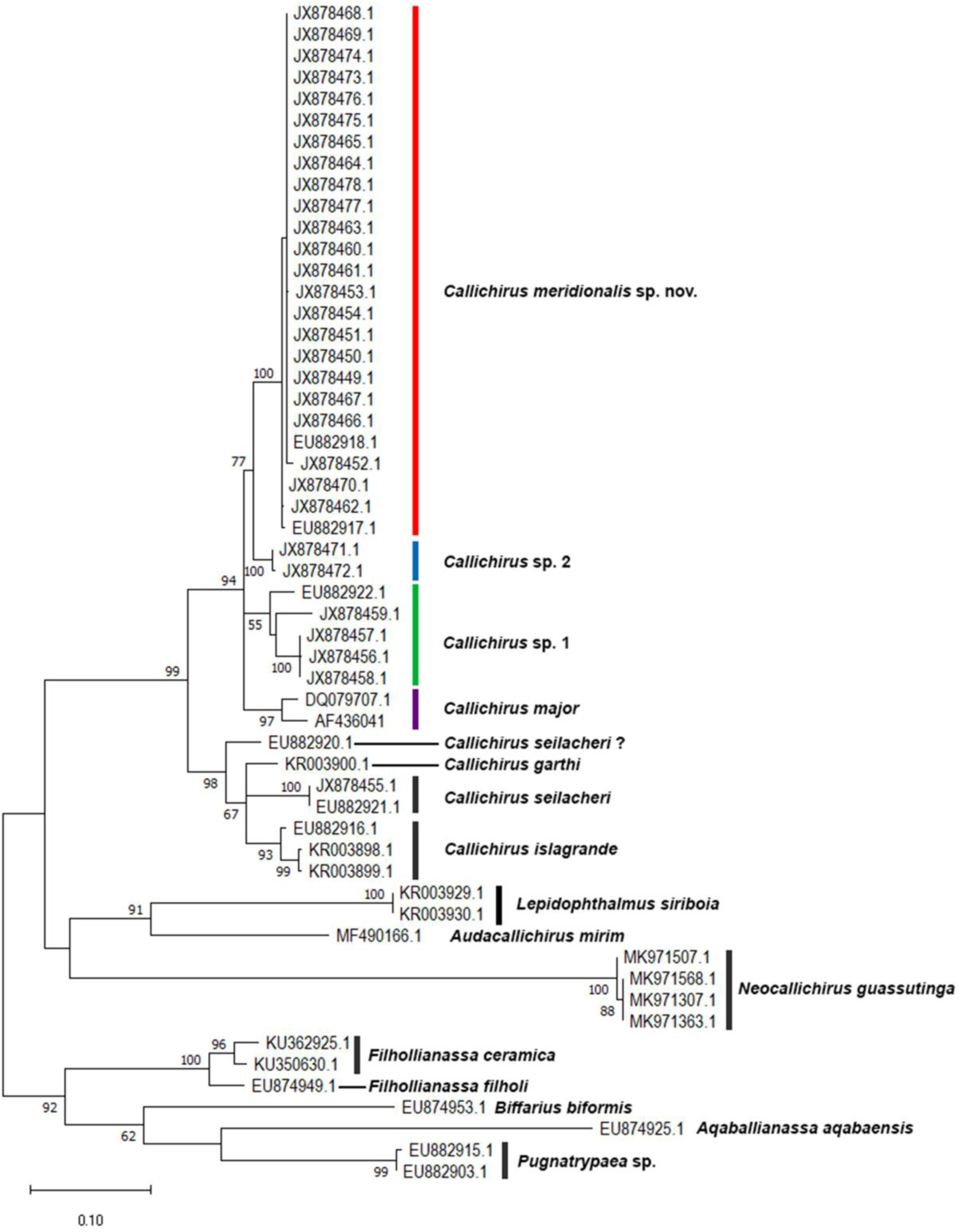
Molecular phylogenetic tree represented as maximum likelihood topology of the partial mitochondrial DNA sequence of the 16S rDNA gene to place *Callichirus meridionalis* sp. nov. Nodal support values represent the frequencies observed using 1000 bootstrap pseudo-replicates. Values below 50% are not represented.

Peiró (2012) corroborated the distinctness of the Brazilian *Callichirus* with molecular data (16S), however, mistakenly assigning the Brazilian specimens to Ortmann’s species in combination with the genus *Callichirus*. Surprisingly, he designated [sic] several males and females from Santos (Brazil) as syntypes [sic] for what he considered to be “*Callichirus macrotelsonis* Ortmann, 1893”. Clearly enough, the fourth larval stage described by Ortmann (1893) is not a *Callichirus* and probably not even a callichirid or callianassid larvae, therefore, Peiró’s action has no taxonomic nor nomenclatural value.

Rio (2018), based on detailed morphological studies, placed the Brazilian *Callichirus* in a new species, *Callichirus brasiliensis* Rio, 2018, but her nomenclatural action in a master’s dissertation is invalid as it does not fulfil the criteria of the ICZN (1999: Articles 8 and 9). This has been corrected in the present paper.

### Neotype designation for Callichirus major (Say, 1818)

Say (1818: 238) described *Callianassa major* (subsequently transferred to *Callichirus* by Stimpson, 1866) from the bay shore of the river St. John, east Florida upon on a single specimen originally housed at the Academy of Sciences of Philadelphia. The holotype has not been found in the collections of the Academy (see Spamer and Bogan 1992: 162). This information has been confirmed by a recent inquiry at the Academy (P. Callomon, personal communication 2020). White (1847: 70) listed 2 “arms” of “*Callianassa major*” presented by Th. Say to the Natural History Museum in London (see Spamer and Bogan 1992: 162). However, no evidence exists to prove that these two “arms” belong to the holotype of *C. major*. Here, the male tl 77 mm (USNM 228086) from Indian River, Flat Just Inside Saint Lucie Inlet, Florida (Manning and Felder 1986) is selected as the neotype of *C. major* (Say, 1818) (Fig. 1a–e) in order to settle the defining characters of *C. major* s. str. and, therefore, ensure the correct use of this name.

### Molecular analysis

The divergence rates between *C. major* s. str. (northern localities) and *C. meridionalis* sp. nov. (southern localities) is 5%, the same value as for the well stablished *C. islagrande* and *C. garthi* and *Filhollianassa filholi* (A. Milne-Edwards, 1879) and *F. ceramica* (Fulton & Grant, 1906) (Table 2). Thus, morphology (see below) and molecular support *C. major* and *C. meridionalis* sp. nov. as distinct species.

*Callichirus* sp. 1 and *Callichirus* sp. 2 (Fig. 5), from the eastern Pacific and Caribbean Colombia, respectively, have a divergence rates of 3% from *C. meridionalis* sp. nov., and 5% from *C. major* (Table 2) These divergence rates also support the separation of *Callichirus* sp. 1 and *Callichirus* sp. 2 into new species. However, additional material of both is needed for morphological studies before conclusions are reached.

The *Callichirus* representatives were found to be nested within a well-supported clade (Fig. 5). The terminals previously assigned to *C. major* s. l. were recovered into a well-supported clade and the sistergroup relationship between *C. meridionalis* sp. nov. and *Callichirus* sp. 2 is also well-supported (bootstrap values of 94 and 77, respectively). *Callichirus* sp. 1 representatives grouped together (bootstrap value 55) but their relationship with *C. major* s. str. and the clade (*C. meridionalis* sp. nov. + *Callichirus* sp. 2) is still unresolved. Resolution of this polytomy will require using additional characters.

### Morphological analysis

Morphologically, the new species differs from *C. major* s. str. by a combination of characters including (Figs. 2b, 3a, 4d *versus* Figs. 6a–f): (i) ocular peduncle reaching to about 2/3 of the length of the first article of the antennular peduncle (Fig. 2b) (*versus* ocular peduncle reaching to the limit between the first and second articles of antennular peduncle, or a little beyond, in *C. major*, Fig. 6a) (see also Manning and Felder 1986: fig. 1a); (ii) second article of the antennular peduncle about two times longer than the distal one (*versus* second article 2.4 times longer than distal one in *C. major*) (see also Manning and Felder 1986: fig. 1a); (iii) male major cheliped dactylus bifid-tipped (Fig. 3a) (*versus* single-tipped male major cheliped dactylus in *C. major*, Fig. 6b) (see also Manning and Felder 1986: fig. 1c); (iv) male major cheliped with fixed finger armed with only one triangular tooth at the midlength of the cutting edge and the dactylus with a large rounded tooth close to articulation with propodus (Fig. 3a) (*versus* fixed finger armed with two teeth, one proximal small triangular tooth and a large rounded tooth at the midlength of the cutting edge, and the dactylus devoid of tooth and, instead, with a large dentate ridge extending from proximal half of the finger to its distal third, being the remaining surface unarmed to the curved tip in *C. major*, Fig. 6b); (v) male major cheliped fixed finger blunt and slightly curved upward (Fig. 3a) (*versus* male major cheliped fixed finger acute and strongly curved upward in *C. major*, Fig. 6e); (vi) male pleopod 2 exopod well developed (Fig. 4d) (*versus* male pleopod 2 exopod reduced in *C. major*, Figs. 6c, d); (vii) first pair of female pereopods, unequal in size and dissimilar in shape (Figs. 3b, d) (*versus* first female pereopods subequal and similar in *C. major*); and (viii) the female major cheliped, with a large meral hook on flexor margin (Fig. 3b) (*versus* that meral hook absent in *C. major*, Fig. 6f).

**Fig. 6.**
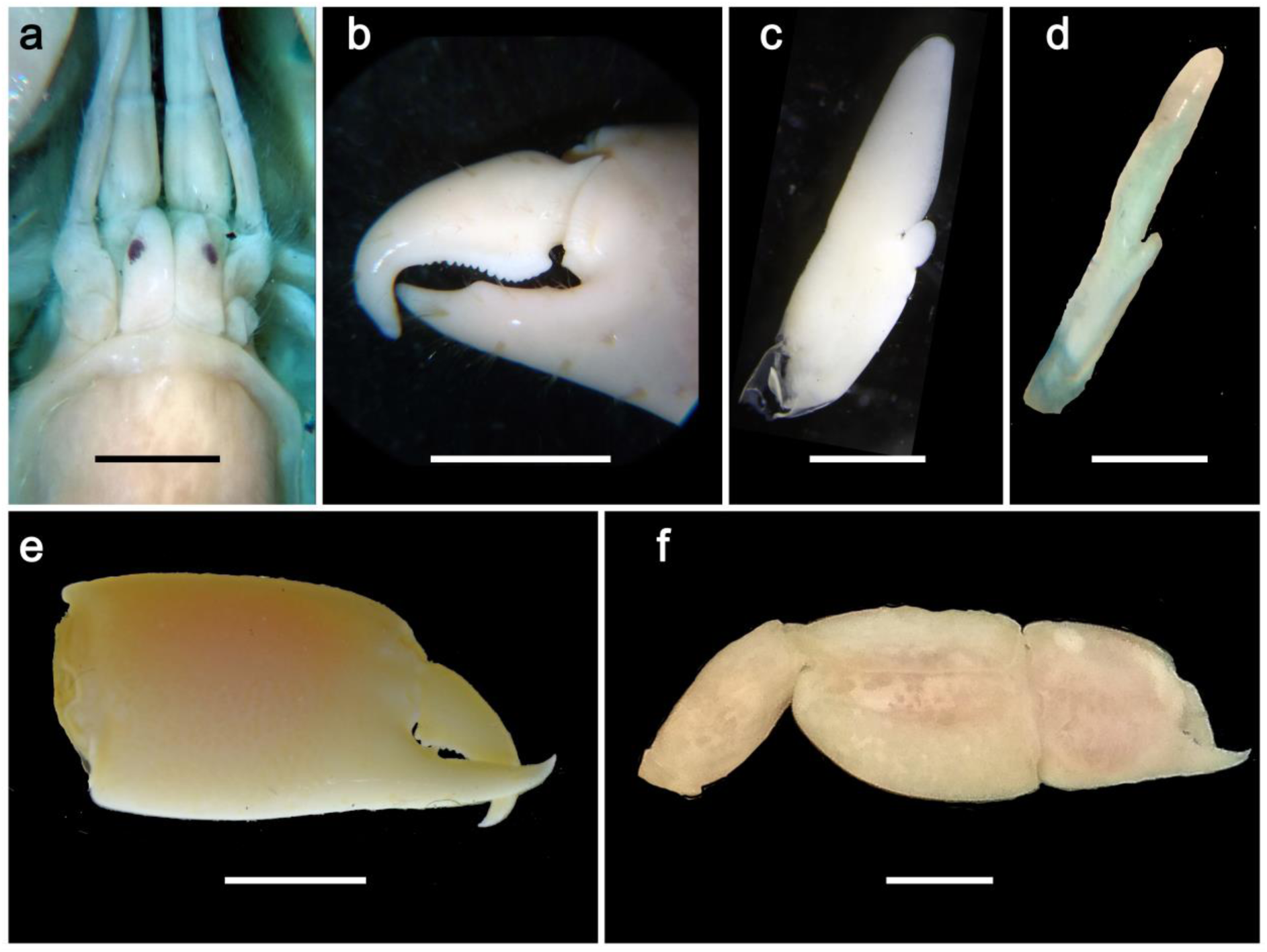
*Callichirus major*, a, b, neotype, male cl 13.2 mm, Indian River, Florida, USA, USNM 228086. c, e, male cl 20.6 mm, New River, North Caroline, USA, USNM 266227. d, male cl 24.4 mm, Louisiana, Gulf of Mexico, USA, USNM 79171. f, female cl 16.3 mm, Indian River, Florida, USA, USNM 228087. a, carapace front, eyestalks, and antennular and antennal peduncles, dorsal view; b, male major cheliped claw, lateral view; c, d, male second pleopod, external surface; e, male major cheliped claw, lateral view; f, female major cheliped, lateral view. Scale bars: a = 2 mm; b, e, f = 1 cm; c, d = 1 mm.

*Callichirus meridionalis* sp. nov. can be easily distinguished from the four other American congeners by the following combination of characters: (i) ocular peduncle elongate with obtuse tip (postcorneal projection absent), not exceeding the second article of the antennular peduncle (*versus* ocular peduncle elongate with postcorneal projections exceeding the second article of the antennular peduncle in *C. garthi*, *C. islagrande* and *C. seilacheri*); (ii) second article of antennular peduncle slightly exceeding the fourth article of antennal peduncle (*versus* second article of antennular peduncle noteworthily exceeding the fourth article of antennal penduncle in *C. garthi* and *C. islagrande*); (iii) maxilliped 3 merus with distal margin strongly oblique (*versus* distal margin slightly oblique in *C. garthi*, *C. islagrande* and *C. seilacheri*); (iv) male major cheliped having the ventral margin of ischium unarmed (*versus* ventral margin of ischium armed with a hook in *C. islagrande* and *C. seilacheri* and a lobe in *C. garthi*); and (v) male pleopod 2 with the endopod well developed and exopod unsegmented (*versus* male pleopod 2 with the endopod reduced and exopod bisegmented in *C. santarosaensis*) (cf. Manning and Felder 1986, figs. 2a-c; Hernáez et al. 2015, figs. 2a,b, 3a,b; Sakai and Türkay 2012, fig. 10h; Hernáez et al. 2018, figs. 2b,c; see also Figs. 2b, h, 3a, 4d).

Sakai and Türkay (2012) claimed that *C. santarosaensis* stands apart from *C. major* in that (i) the antennular peduncle is about as long as the antennal peduncle and (ii) the telson is bisectioned, slightly converging toward the truncate end. However, Felder and Dworschak (2015) have convincingly showed that misinterpretations of the morphology of antennula, antenna, and telson led Sakai and Türkay (2012) astray, so that no morphological characters remain that justify *C. santarosaensis*. Yet, Felder and Dworschak (2015) argued that the name *C. santarosaensis* can still be applied “on the basis of geographical origins of materials, underpinned by genetic analyses (*sensu* Staton and Felder 1995) when possible”. However, Sakai and Türkay (2012), but also Felder and Dworschak (2015), overlooked that the male second pleopod endopod is greatly reduced in *C. santarosaensis* (Sakai and Turkay 2012: 746, fig. 10h) while well developed in *C. major* s. str. (Figs. 6c, d). This character can be used with confidence in separating *C. santarosaensis* from *C. major* s. str., unless Sakai and Türkay (2012) have confused the endopod with the exopod in *C. santarosaensis*.

*Callichirus meridionalis* sp. nov. stands apart from *C. adamas*, the only non-American species in the genus, in the presence of ocular peduncles with obtuse tips without elongated postcorneal projections (*versus* ocular peduncles with elongated postcorneal projections in *C. adamas*) and by the male second pleopod with well develop exopod (*versus* male second pleopod exopod vestigial or strongly reduced in *C. adamas*) (Kensley 1974, figs. 1a, 2h; see also Figs. 2b, 4d).

*Callichirus* now consists of 7 species, namely: *C. adamas* (known from Senegal to Orange River mouth, South Africa), *C. garthi* (known from Huanchaco, Peru to Tubul, Chile), *C. islagrande* (currently known from the Gulf of Mexico), *C. meridionalis* sp. nov. (from Pará to Santa Catarina, Brazil), *C. major* (from North Caroline, USA to the Caribbean coast of Colombia), *C. santarosaensis* (known only from the northern coast of the Gulf of Mexico, Florida, USA) and *C. seilacheri* (From Jalisco, Mexico to Puntarenas, Costa Rica).

*Callichirus meridionalis* sp. nov. is the fourth western Atlantic species in *Callichirus*. However, the specimens from the Caribbean Colombia probably belong to an undescribed species sister to *C meridionalis* sp. nov. Both differ from *C. major* s. str. in having shorter ocular peduncles that do not reach to the second article of antennular peduncle, and male pleopod 2 with well-developed exopod. However, the Caribbean Colombia population and *C. meridionalis* sp. nov. can be easily separated from each other in that the Colombian specimens have single-tipped male major cheliped dactylus, whereas *C. meridionalis* sp. nov. has bifid-tipped male major cheliped dactylus, and in the lack of a hook on the ventral margin ventral margin of merus of the female major cheliped, whereas the ventral margin of the female major cheliped merus is armed with a hook in *C. meridionalis* sp. nov. The following key facilitates the identification of the species of *Callichirus*.

## Key to the species of the genus *Callichirus* Stimpson, 1866

1.- Ocular peduncle elongate with obtuse tip, not exceeding or slightly exceeding to the junction between the first and second articles of the antennular peduncle……………2

- Ocular peduncle elongate with postcorneal projection noteworthy exceeding to the junction between the first and second articles of the antennular peduncle……………5

2.- Ocular peduncle reaching to about 2/3 of the length of the first article of the antennular peduncle. Male pleopod 2 with endopod and exopod well developed……………3

- Ocular peduncle reaching to the limit between the first and second articles of antennular peduncle, or a little beyond. Male pleopod 2 with endopod or exopod reduced……………4

3.- Male major cheliped dactylus with bifid-tipped. Female major cheliped merus armed with ventral hook………………………….…………………….. *C. meridionalis* sp. nov.

- Male major cheliped dactylus with single-tipped. Female major cheliped merus unarmed………………………………………….*Callichirus* sp. 2 (Caribbean Colombia)

4.- Male pleopod 2 with endopod underdeveloped. Antennular peduncle about as long as antennal peduncle…………………………………………………………………… *C. santarosaensis*

- Male pleopod 2 with exopod reduced. Antennular peduncle longer than antennal peduncle………………………………………………………………………………………… *C. major* s. str.

5.- A1 peduncle article 2 reaching as far as A2 peduncle article 4. Maxilliped 3 propodus with V-shaped depression distally……………………………….. *C. seilacheri*

- A1 peduncle article 2 exceeding A2 peduncle article 4. Maxilliped 3 propodus rounded or straight distally……………6

6.- Male major cheliped ischium unarmed ventrally. Telson emarginate posteriorly, with rounded tooth on distomedial depression………………………………………. *C. garthi*

- Male major cheliped ischium armed with a ventral hook. Telson emarginate posteriorly, unarmed distomedial depression……………7

7.- Male pleopod 2 with exopod reduced. Male major cheliped merus unarmed ventrally……………………………………………………………………….. *C. adamas*

- Male pleopod 2 with exopod well developed. Male major cheliped merus armed with ventral hook………………………………………………………………… *C. islagrande*

## Acknowledgements

Firstly, we would like to thank very much our friends Marcos Tavares and William Santana for their help in the construction, organization, analysis and conclusions of this work. Without their help and work we would not have been able to bring this work to fruition. Also, we are grateful to Rafael Lemaitre and Karen Reed, who kindly provided access to material of *Callichirus* and provided working space at the USNM; Paul Callomon (ASP, Academy of Sciences of Philadelphia) and Paul F. Clark and Miranda Lowe (NHM, Natural History Museum, Lodon), who kindly searched for the holotype o *Callichirus* major in the collections of the ASP and NHM, respectively; Joana D’Arc de Jesus Pinto and Maria José de Souza Coelho (MZUSP) for the help with the MZUSP collections. We also thank Jessica Colavite (UNESP) for the help during the molecular analyses.

## Funding information

PH and MSM thank FAPESP (Fundação de Amparo à Pesquisa do Estado de São Paulo) for financial aid through a post-doctoral grant (process 2015/09020-0) and a PhD grant (process 2018/17718-5), respectively. JPPR thanks CAPES (Coordenação de Aperfeiçoamento de Pessoal de Nível Superior) for financial aid through a MSc grant (process 1688290). MAAP and MT thank CNPq (Conselho Nacional de Desenvolvimento Científico e Tecnológico) for financial aid through research grants 305714/2015-5 and 303122/2016-1, respectively.

## Compliance with ethical standards

### Conflict of interest

The authors declare that they have no conflict of interest.

### Ethical approval

All applicable international, national, and/or institutional guidelines for the care and use of animals were followed by the authors.

### Sampling and field studies

All necessary permits for sampling and observational field studies have been obtained by the authors from the competent authorities. Licenses to collect zoological material issue to PH (#51578-1, #58845-1) were provided by Brazilian Institute of the Environment and Renewable Natural Resources (SISBIO/IBAMA-MMA).

### Data availability

All data generated or analyzed during this study are included in this published article.

## Notes

### Competing Interest Statement

We have conflict of interest with the following colleagues:
Dr. Fernando L. Mantelatto
Dr. Rafael R. Robles
Dr. Douglas F. Peiro
Dr. Daryl Felder
Dr. Peter Dworschak
Dr. Ingo S. Wehrtmann
Dr. Enrique MacPherson

